# An enhanced transcription factor repressilator that buffers stochasticity and entrains to an erratic external circadian signal

**DOI:** 10.1101/2022.10.10.511622

**Authors:** Steven A. Frank

## Abstract

How do cellular regulatory networks solve the challenges of life? This article presents computer software to study that question, focusing on how transcription factor networks transform internal and external inputs into cellular response outputs. The example challenge concerns maintaining a circadian rhythm of molecular concentrations. The system must buffer intrinsic stochastic fluctuations in molecular concentrations and entrain to an external circadian signal that appears and disappears randomly. The software optimizes a stochastic differential equation of transcription factor protein dynamics and the associated mRNAs that produce those transcription factors. The cellular network takes as inputs the concentrations of the transcription factors and produces as outputs the transcription rates of the mRNAs that make the transcription factors. An artificial neural network encodes the cellular input-output function, allowing efficient search for solutions to the complex stochastic challenge. Several good solutions are discovered, measured by the probability distribution for the tracking deviation between the stochastic cellular circadian trajectory and the deterministic external circadian pattern. The solutions differ significantly from each other, showing that overparameterized cellular networks may solve a given challenge in a variety of ways. The computation method provides a major advance in its ability to find transcription factor network dynamics than can solve environmental challenges. The article concludes by drawing an analogy between overparameterized cellular networks and the dense and deeply connected overparameterized artificial neural networks that have succeeded so well in deep learning. Understanding how overparameterized networks solve challenges may provide insight into the evolutionary design of cellular regulation.

## Introduction

Transcription factor (TF) networks control cellular response. The concentrations of the TF proteins feed into the TF network, which then stimulates or represses the expression of various genes. The TF network computes an input-output function: TF concentrations in, gene expression out.

A key puzzle of cellular design concerns how TF input-output functions solve the challenges of life. The puzzle poses two questions. First, what functional relation between inputs and outputs solves the biological challenge? Second, how is the functional relation encoded molecularly?

This article focuses on the first problem, the functional solution to the biological challenge. Following a prior article, the challenge concerns maintaining a circadian rhythm of molecular concentrations (Frank, 2022c). The rhythm must buffer stochastic fluctuations in molecular concentrations.

An internal rhythm perturbed by stochasticity cannot retain perfect periodicity without an external signal. In this case, an external circadian signal appears and disappears randomly. That external signal provides an occasional opportunity for entrainment. However, the erratic external signal does not provide sufficient information for the cell to achieve good periodicity by simply mirroring the signal. Instead, the TF network must entrain to the external signal when available and otherwise buffer internal molecular stochasticity to maintain its internal rhythm in the absence of external information.

I use an artificial neural network to search for a TF input-output function that solves this circadian challenge. I embed the TF network in a stochastic differential equation that tracks the concentrations of mRNAs transcribed by gene expression and the associated TF proteins translated from the mRNAs. The TF network takes the TF protein concentrations as inputs and produces as outputs the mRNA transcription rates for the various TF genes.

I studied the same circadian challenge in the prior article (Frank, 2022c). That prior article tried to solve the functional and mechanistic parts of the puzzle simultaneously. Mechanistically, I encoded the TF input-output function using the currently favored thermodynamic model for TF binding to DNA gene promoter regions and the consequences of that binding for mRNA transcription rates (Bintu, Buchler, Garcia, Gerland, Hwa, Kondev & Phillips, 2005; Bintu, Buchler, Garcia, Gerland, Hwa, Kondev, Kuhlman, et al., 2005).

Among many computer runs in that prior study, I found only one solution that provided reasonably good circadian tracking and entrainment based on explicit thermodynamic parameters for the TF network. That solution was very difficult to find computationally and was sensitive to changes in the thermodynamic parameters. Additionally, although the thermodynamic model I used is the most widely favored description for the molecular mechanism, it is very unlikely for that model to be an accurate and complete description of the actual molecular mechanism.

This article gains by focusing solely on the functional challenge of mapping inputs to outputs without concern for the mechanism. In particular, I use an artificial neural network to encode the functional mapping instead of the TF thermodynamic model. Once we have an understanding of the kinds of functional relations that solve the challenge, we can then search for candidate molecular mechanisms that could potentially encode those functional relations.

For example, given a functional solution encoded by a neural network, one could then fit the classic thermodynamic model to that functional solution. That fitting of the molecular mechanism to the functional solution would be useful when attempting to engineer the mechanism in actual cells and measure their molecular dynamics.

Prior modeling approaches searched for TF networks to achieve particular dynamics (Bolouri & Davidson, 2002; Kauffman et al., 2003; De Jong et al., 2004; Mishra et al., 2018; Patel & Bush, 2021). The approach in this article has several technical advantages. Neural networks are often ideal function approximators (Goodfellow et al., 2016). Computationally, they scale very well to higher dimensions, can easily use automatic differentiation for optimization, and can be embedded within stochastic differential equations while maintaining the great computational advantages of automatic differentiation (Baydin et al., 2018; Rackauckas et al., 2020; Frank, 2022b). Without these technical advantages, computational optimization of TF networks is difficult and has not previously been studied in a widely applicable way. The computer code provided with this article can easily be adapted to study other biological challenges. Additional studies will eventually give a sense of the kinds of input-output mappings required of TF networks to solve the demands of life.

As future studies accumulate, we may find that the evolutionary success of TF networks and the computational success of neural networks arise from a common foundation. Both networks may induce essentially the same geometric manifold of evolutionary or learning dynamics on which improving performance plays out (Frank, 2017).

## Methods

This article extends the approach in Frank (2022c). The most important change replaces the prior thermodynamic model for the TF input-output response function with a computational neural network, which leads to greatly improved optimization outcomes.

I wrote the computer code in the Julia programming language (Bezanson et al., 2017). I used the SciML packages to optimize differential equations (Rackauckas et al., 2020). Efficient optimization depends on automatic differentiation (Baydin et al., 2018; Margossian, 2019), which is built into the SciML system. The source code for this article provides the details for all calculations and plotting of results (Frank, 2022a).

The goal is to optimize a TF regulatory control system in order to track an environmental target. I first describe the TF system and then describe the environmental target.

The deterministic component of the TF system dynamics is given by the temporal derivatives for numbers of mRNA molecules, *x*, and the TFs produced by those mRNAs, *y*, as

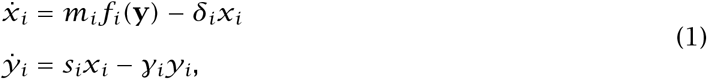

for *m*_*i*_ as the maximum mRNA production rate, *δ*_*i*_ as the mRNA decay rate, *s*_*i*_ as the TF production rate per mRNA, and *γ*_*i*_ as the decay rate of the *i =* 1, …, *n* TFs (Marbach et al., 2010).

The function *f*_*i*_ transforms the numbers of TFs in the vector **y** into the production level of the *i*th mRNA, varying between 0 for complete repression and 1 for maximum production. In this article, each *f*_*i*_ is a separate neural network that takes *n* inputs as log(**y**).

The neural network architecture follows a standard general form, with the following details.

The first layer of the network has 5*n* output nodes. Each of those nodes sums an affine transformation of each input, *z*, of the form *α + βz*, in which each of those 5*n*^2^ transformations has its own *α* and *β* parameters. The value of each output node for this first layer is transformed by the mish activation function (Misra, 2019)

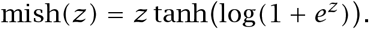

The 5*n* outputs from the first layer form the inputs for another neural network layer, which leads to the final single-valued output that controls the mRNA transcription rate for the associated TF gene. That output arises by first summing affine transformations from all 5*n* inputs and then applying the sigmoid function to that sum, in which the sigmoid function outputs values between 0 and 1, as

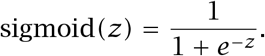

The same architecture is repeated for each of the *n* TFs, with the single-valued output of each network influencing the transcription rate of the associated TF gene.

I added stochastic perturbations to the molecular abundances in eqn 1, making the system a set of stochastic differential equations. Stochastic fluctuations for a molecule with abundance *z* follow 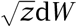 in which *W* is a standard normal variate with mean 0 and standard deviation 1, thus 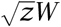 has a standard deviation of 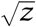

The normalized magnitude of the fluctuations is the ratio of the standard deviation relative to the current abundance, 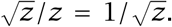. As *z* drops, the normalized fluctuations increase. To prevent fluctuations from becoming too large relative to the abundance, which can cause negative abundance values in the numerical analysis, for *z* ≤ 16, I replaced 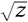 with *z/*4. Thus, we may write the stochastic component as 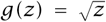 for *z >* 16 and *z/*4 for *z* ≤ 16, leading to the stochastic differential equations

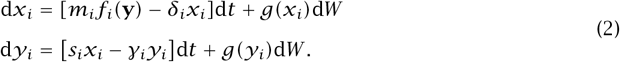

I calculated the trajectories of these stochastic differential equations with the Julia DifferentialEquations.jl package (Rackauckas & Nie, 2017), using the Ito implicit method ISSEM. The Julia SciML libraries optimize the performance of the TF system by passing the automatic differentiation procedure through the stochastic differential equations, including the embedded neural networks.

The goal is for the stochastic TF system to maintain a circadian rhythm given only a sporadically present external circadian signal, as described in Frank (2022c) and summarized here. In particular, the design goal is for TF 1 *(y*_1_) to follow a 24h period. TF abundance above *S =* 10^3^molecules per cell corresponds to an “on” state for daytime. Below that threshold, the cell is in an “off” nighttime state.

The optimization procedure seeks a parameter combination that minimizes the loss measured as the distance between the target circadian pattern shown in the gold curve and the transformed abundance of TF 1 in the green curve in Fig. 1a. In particular, the daily target rhythm follows

**Figure 1:**
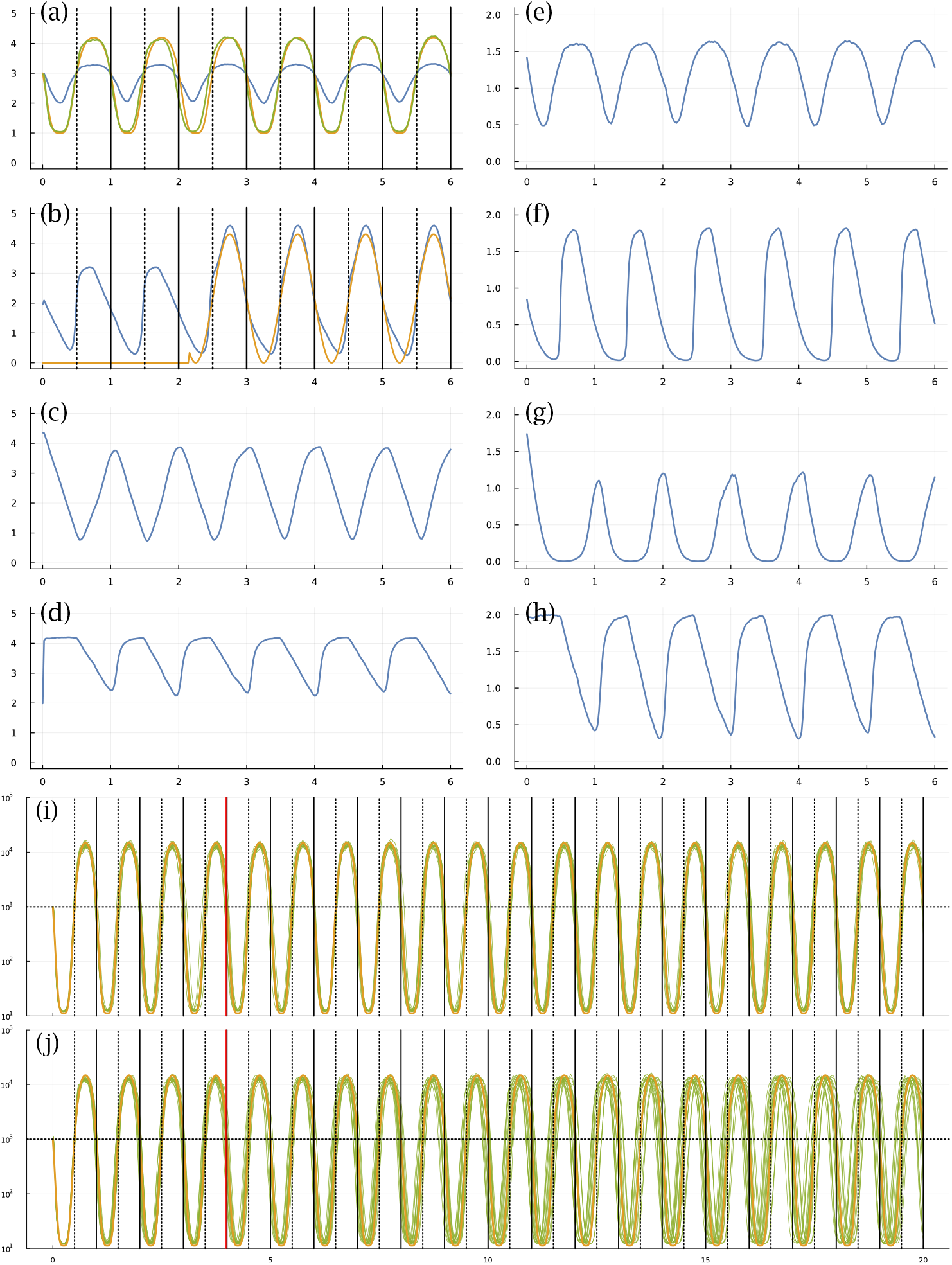
Circadian dynamics of the TF network from run sde–4_1_t4. Panels (a-d) show the stochastic dynamics of the TF proteins. The dynamics of the matching mRNAs that produce the proteins are shown in panels (e-h). The time period is six days, along the *x*-axis of each of these panels. The vertical lines in (a,b) show entry into daylight (dotted) and nighttime (solid). The *y*-axis is log 10(1 *+ y)* for number of molecules per cell, *y*. In (a), the optimization procedure attempts to match the number of TF 1 molecules (blue curve) to a circadian rhythm by minimizing a loss value. To calculate the loss value, begin with the number of TF 1 molecules, *y*, transformed by a Hill function, 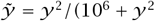, to yield the green curve, which traces 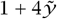. The gold curve traces the target circadian pattern. The loss value to be minimized is the sum of the squared deviations between the gold and green curves at 50 equally spaced time points per day. The number of TF 2 proteins in (b) is influenced by the internal cellular dynamics and is also increased in response to an external daylight signal that switches on and off randomly (Frank, 2022c). It is initially off. The average waiting time for a random switch in the presence or absence of the signal is *w =* 2, measured in days. In this example, the signal turns on during the night of the third day and stays on for the remaining days shown. Because the switching is random, daylight can be present or absent for several days in a row, or it can switch on and off several times in one day. In panel (a), stochastic molecular perturbations push the cellular rhythm (green curve) behind the actual circadian pattern (gold curve) during the first few days in this particular realization of the stochastic dynamics. When the daylight signal appears in day 3, the system entrains to the external signal, closely matching the target circadian pattern for the remaining days shown. Panels (i,j) are described in the text. See Frank (2022c) for additional details.

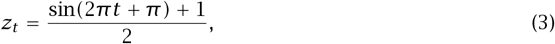

with *t* in days and passage across 0.5 corresponding to transitions between day and night. The loss is

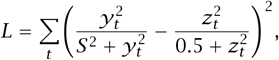

in which *y*_*t*_ is the abundance of TF 1, and the transformations of *y* and *z* are Hill functions that normalize values to make the different scales comparable, and *S =* 10^3^ is the cellular state switch rate given above. For the summation, the time values start at *t =* 0 and increment by 0.02 until the end of time for a particular analysis. The values for *y* and *z* in Fig. 1a are normalized to the scaling of the plots, as described in the figure legend.

A stochastic system inevitably diverges from the target circadian trajectory. In this model, the system may use an external daylight entrainment signal to correct deviations. To pass information about the external signal into the system, I added a strong boost to the production rate of TF 2 *(y*_2_) in proportion to the intensity of external light. In particular, the rate of change in TF 2 abundance in the presence of the external light signal is augmented by 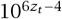, in which *z*_*t*_ is the daily rhythm in eqn 3.

Initially, the light signal is absent. The signal switches on and off randomly. In Fig. 1b, the gold curve shows the external light signal, which switches on in the third day and stays on for theremaining days shown. The blue curve traces the abundance of TF 2. Thus, the overall challenge is for TF 1 to track the circadian pattern, given internal stochastic molecular dynamics and a randomly occurring external entrainment signal that influences the abundance of TF 2.

## Results

### Overview

Computational optimization finds parameters for eqn 1 that influence the stochastic dynamics of mRNAs and TF proteins. Different runs of the optimization procedure converged to different parameters. Figure 1 illustrates the results for one run.

In that figure, panel (a) shows the target circadian pattern in gold and the system’s circadian pattern in green. The optimization goal is to minimize the total squared distance between those two curves (Frank, 2022c). The blue curve in that panel shows the molecular abundance of the TF 1 protein. The green curve arises by transforming the blue curve by a Hill function, as explained in the figure caption. The *x*-axis shows time measured in days.

The optimization begins by attempting to fit the parameters to the circadian pattern for part of the first day. Then, the algorithm slowly increases the temporal range in a sequence of steps, for each step allowing the fit to adjust to the additional time range. The final time range for fitting the parameters in these runs was four days. Once the fit is obtained, I studied the match between the target circadian pattern and the system dynamics over a time period greater than the four days used for fitting.

I wrote the code to emphasize the chance of finding a good fit rather than to maximize the speed of the fitting process. A single optimization run takes about ten days when using 12 computational threads on a 2022 Apple Studio Ultra M1 computer. I analyzed eight full runs for this study, which was sufficient to find several good fits and to illustrate a range of outcomes.

### Dynamics

In Fig. 1, panel (b) shows the dynamics of the TF 2 protein. The blue curve is the abundance of that TF. The gold curve shows the external circadian entrainment signal, which may be interpreted as the external light intensity experienced by the cell. The entrainment signal comes and goes randomly. For these runs, the signal begins in the off state and then switches on and off randomly with a waiting time drawn from an exponential distribution with a mean of two days. The TF 2 protein increases in abundance in response to the light intensity (Frank, 2022c).

Overall, panels (a-d) show the dynamics of the four TF proteins. For each TF protein, the matching panel to the right shows the dynamics of the mRNAs that encode the TFs.

The gold curve in panel (i) shows the target circadian pattern over 20 days. Each of the 20 green curves shows the cellular circadian state for a run of the stochastic dynamics. As in panel (a), cellular state is calculated as a Hill transformation of the abundance of TF 1. For each run in panel (i), the external entrainment signal comes on and off randomly with a mean waiting time of two days, as in panel (b).

Panel (j) repeats the setup in panel (i), except that the external signal comes on and off randomly with a mean waiting time of 1000 days. Because the signal starts in the off state, almost always the signal never comes on. The green curves therefore show the ability of the system to maintain its circadian rhythm in the absence of any external signal. Inevitably, the stochastic fluctuations of molecular concentrations cause the cellular rhythm to diverge from the target pattern.

Figures 2 and 3 show the same data for two other optimization runs. The following subsections compare the different optimization runs. For all runs, the settings, optimized parameters, output values, and code for making the figures are included in the online repositories listed in the Data Availability statement at the end of the article.

**Figure 2:**
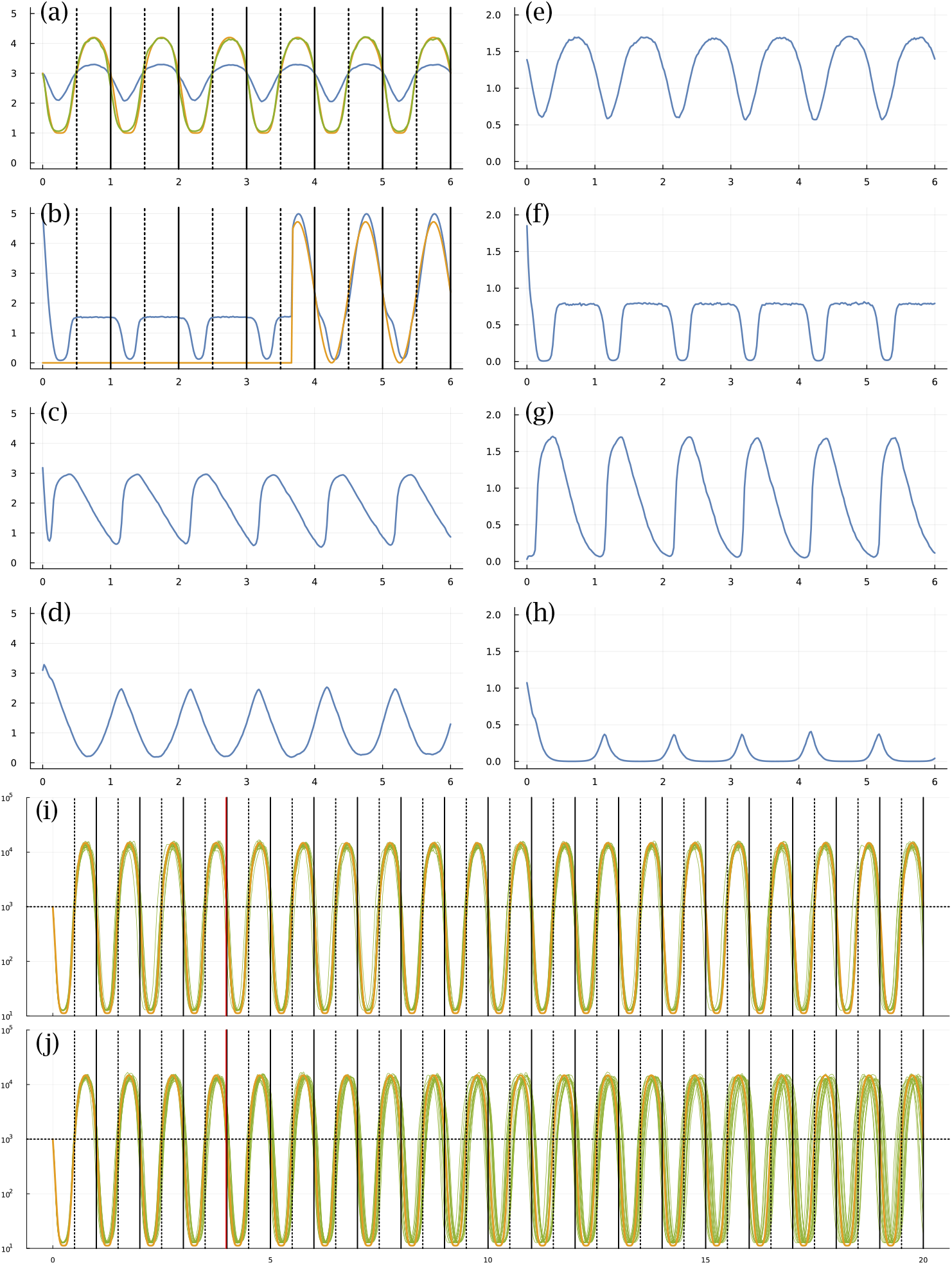
Circadian dynamics for run sde–4_2_t4.

**Figure 3:**
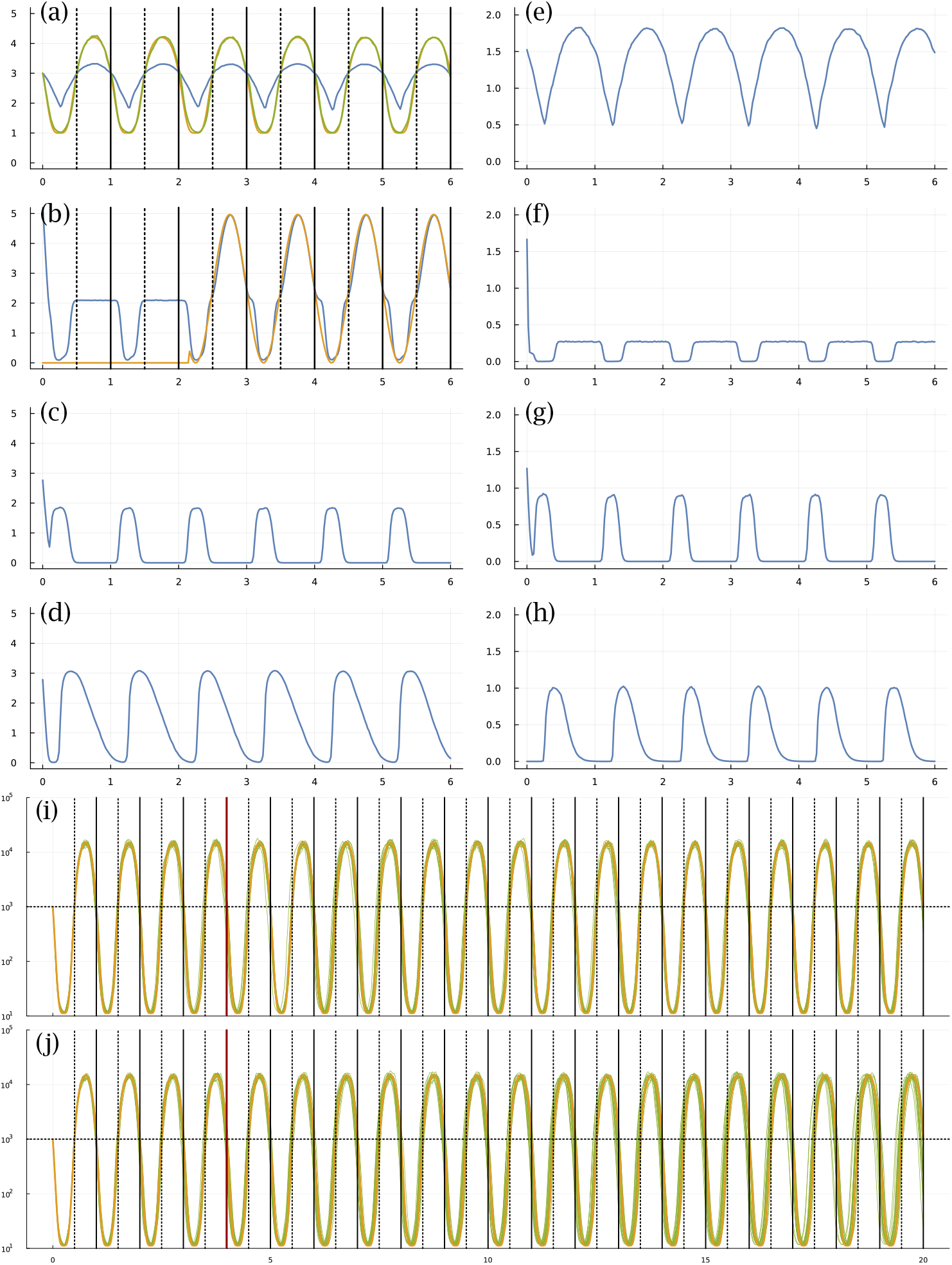
Circadian dynamics for run sde–4_8_t4.

### Deviation from the target pattern

Figure 4a shows the deviations of each optimized system from the target circadian pattern. For a single realization, such as a green curve in panel (i) or (j) of the previous figures, I calculated the deviations as follows. The entry into external daytime occurs at the vertical dotted line in each daily interval, and the gold curve crosses the horizontal dotted line at 10^3^. For a cell, the entry into its internal daytime state occurs as the green curve crosses above the horizontal dotted line at abundance 10^3^.

**Figure 4:**
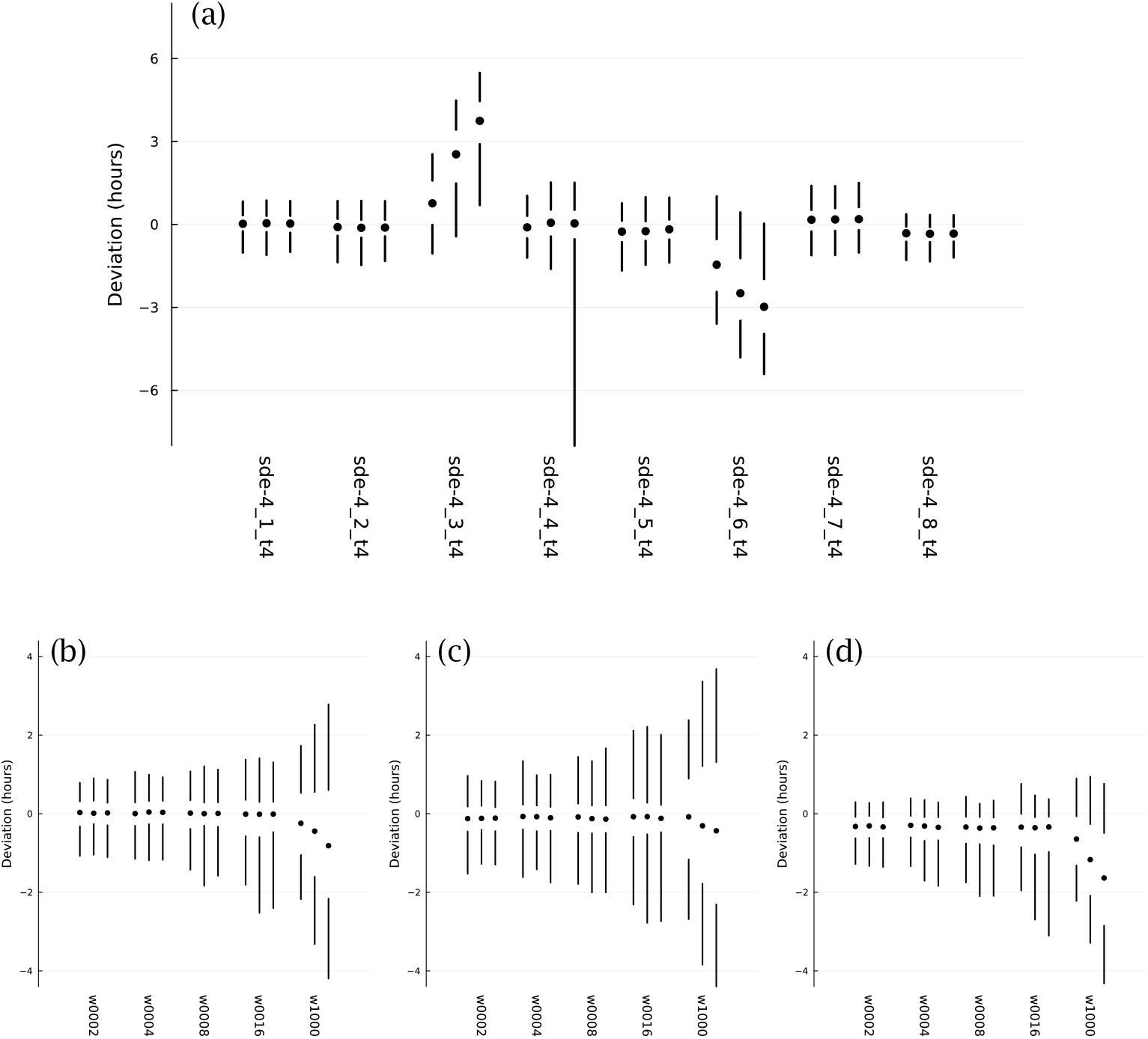
Probability distributions for deviations between the actual circadian pattern of the TF 1 protein and the target circadian pattern. Each set of two vertical lines and a circle describe a distribution, with the 5th to 25th percentiles of the distribution along the lower line, the 50th percentile at the circle, and the 75th to 95th percentiles along the upper line. (a) The overall patterns for the eight independent optimization runs, with the label for each run along the *x*-axis. (b) The distributions for sde–4_1_t4, for different values of *w*, the average waiting time for switching of the external circadian light signal. (c) The distributions for sde–4_2_t4. (d) The distributions for sde–4_8_t4. Further details in the text.

For each day, the cellular deviation is the horizontal distance along the dotted line between the green curve and the gold curve. When the green curve is to the left of the gold curve, the deviation is negative. When it is to the right, the deviation is positive. I transformed the deviation into hours, the daily deviation amount divided by 24. For a realization of the dynamics over *d* days, there are *d* deviation measures for that realization.

For realizations that ran over *d* days, I repeated the measurements over 1000 samples for a total of 1000*d* deviation measures. I then calculated percentiles of the deviations. Looking at the far left of Fig. 4a, the two vertical lines and the circle show those percentiles. The lower line ranges from the 5th to the 25th percentile. The circle marks the 50th percentile. The upper line ranges from the 75th to the 95th percentile.

Thus, each vertical set summarizes the distribution of deviations for a sample of 1000 realizations over *d* days. In the figure, the vertical summary for each distribution occurs within a set of three such distributions. For each set, the left, middle, and right distributions correspond to *d =* 10, 20, 30, respectively, showing the distributions of deviations when realizations run over different numbers of days.

The eight labels below the sets in Fig. 4a provide the names for the optimization runs. I chose for further analysis the two sets on the left and the set on the right. Those three optimization runs had the narrowest distributions for the deviations, in which smaller deviations correspond to better tracking of the target circadian pattern.

The leftmost run, sde–4_1_t4, corresponds to the dynamics in Fig. 1 and to the more detailed summary of deviations in Fig. 4b. The deviation details in Fig. 4b show the distributions induced by different random switching times for the external entrainment signal. Each set of three distributions again corresponds to runs with *d =* 10, 20, 30 days. The different sets correspond to mean waiting times for random exponential switching of 2, 4, 8, 16, and 1000 days.

The mean time of 2 days corresponds to the value used in during optimization and is the default in all other plots unless otherwise noted. The longer mean times illustrate how well the system can hold its internal circadian rhythm when receiving an external entrainment signal less reliably. When the mean is 1000 days, the system almost never receives an external signal. The bigger deviations in that case show the greater difficulty of maintaining a match to the target pattern without the opportunity for entrainment. Although the spreads are greater for larger *d*, the system nonetheless typically keeps a very good circadian pattern for 10 days and a reasonably good pattern for as along as 30 days.

Figure 4c corresponds to sde–4_2_t4, and Fig. 4d corresponds to sde–4_8_t4. The sets do not match exactly between panel (a), which has a mean waiting time of 2 days, and the leftmost sets in panels (b-d), which also have mean waiting times of 2 days. The small mismatches arise because the different panels used different samples of the stochastic trajectories to calculate the distributions.

### TF network input-output functions

Figure 5 shows the TF network input-output relations associated with sde–4_1_t4. Each quadrant presents the output as the relative mRNA production rate for a TF gene. For example, the upper-left block describes the output for TF 1. In each plot, the output level varies from off, in dark orange, to maximally on, in dark purple.

**Figure 5:**
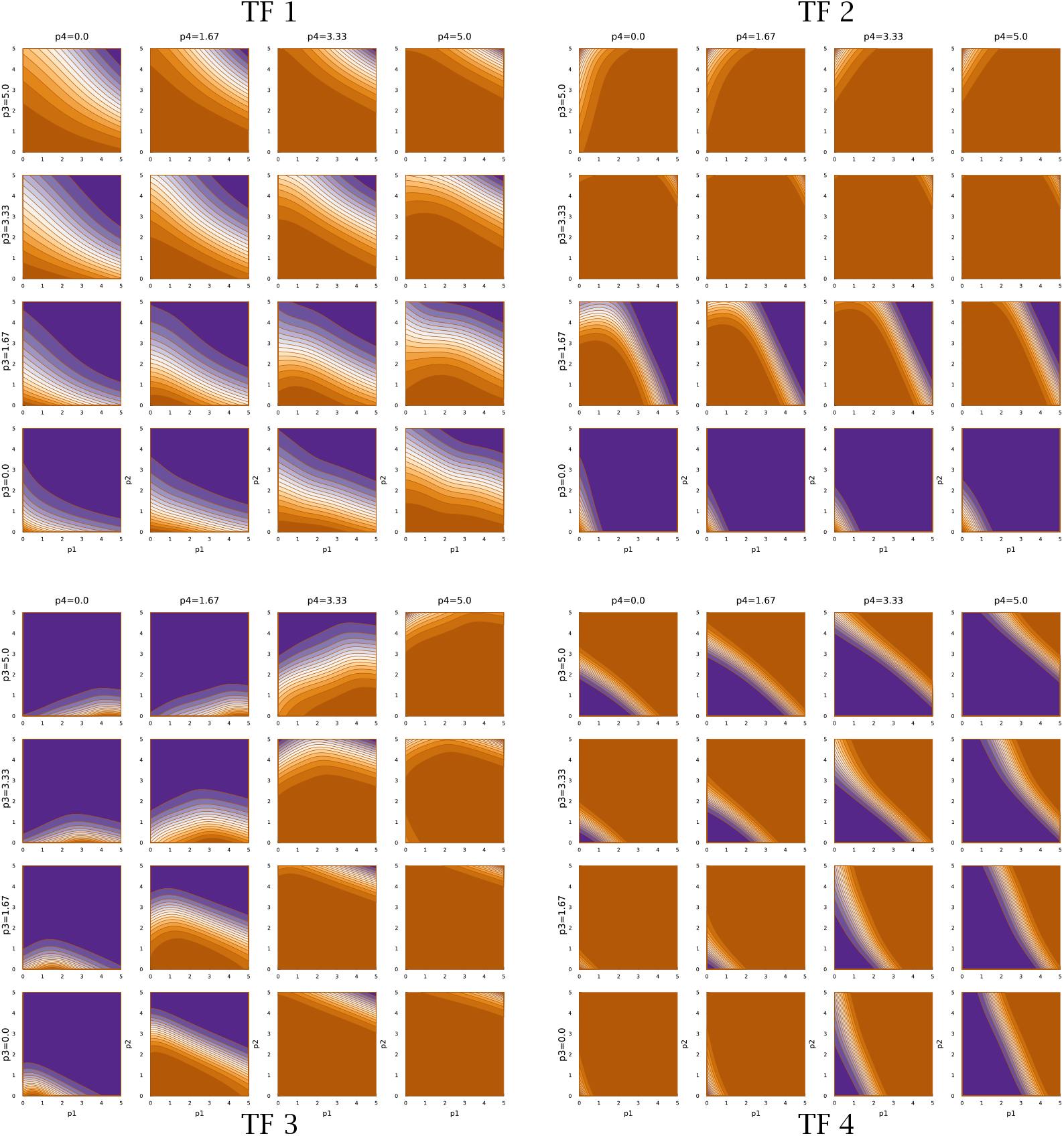
TF network input-output function for run sde–4_1_t4.

The output level depends on four inputs, the abundances of the four TF proteins. The *x* and *y* axes of a plot correspond to the abundances of TF 1 and TF 2, respectively, on a log 10(1 *+ z)* scale for abundance *z*. The abundance of TF 3 rises across the rows of plots, from zero at the bottom of a quadrant to the maximum level at the top. The abundance of TF 4 rises across the columns, from zero at the left to maximal at the right.

The TF network logic can be read from the figure. Consider, for example, the TF 2 block in the upper right that shows the TF 2 mRNA output rate. An increase in TF 3 abundance, going up the rows, associates with repression of TF 2, because increasing orange means lower output. In the TF 3 block (lower left), an increase in TF 4, going right across columns, associates with repression of TF 3. In the TF 4 block (lower right), an increase in TF 2, going right along the *x*-axis of each plot, associates with repression of TF 4.

Thus, TF 3 represses TF 2, and TF 4 represses TF 3, and TF 2 represses TF 4. That cyclic repression defines the basic repressilator circuit, which can cause periodic oscillation of the associated TF proteins (Elowitz & Leibler, 2000). However, in this case, that core repressilator is embedded within a more complex network of modulating inputs. For example, TF 2 output is stimulated by TF 1 when both TF 2 and TF 3 abundances are low.

In this optimization problem, one major function of the TF network is the entrainment to an external circadian signal when available, illustrated by panels (a) and (b) of Fig. 1. Returning to the upper-left quadrant of Fig. 5, note that increasing TF 2 abundance (*y*-axis of plots) stimulates TF 1 expression. Thus, when an external light signal increases TF 2, that rise in TF 2 stimulates a rise in TF 1 expression. A decline TF 2 abundance during the night represses TF 1 expression.

Other inputs modulate the relation between TF 2 and TF 1. For example, high TF 3 and TF 4 prevent TF 1 expression, unless both TF 1 and TF 2 are very high. TF 1 stimulates its own expression on the way up and represses its own expression on the way down.

The second major function of the TF network is to maintain the circadian pattern by buffering the intrinsic biochemical stochasticity. That buffering arises from the various interactions of the TF network logic. One aspect may be the tendency to maintain both TF 1 and TF 2 cycling in synchrony with the target rhythm. Joint rhythmicity provides a partially redundant cycle that can be used to smooth out fluctuations in either TF protein.

Figures 6 and 7 show the TF network logic for sde–4_2_t4 and sde–4_8_t4, respectively. Looking at Fig. 6, one can see that the quantitative relations differ significantly from Fig. 5. For example, most effects of changing abundance in Fig. 5 are monotonic. By contrast, in Fig. 6, the upper-left and lower-right quadrants show reversals in the direction of change in outputs with changes in particular inputs.

**Figure 6:**
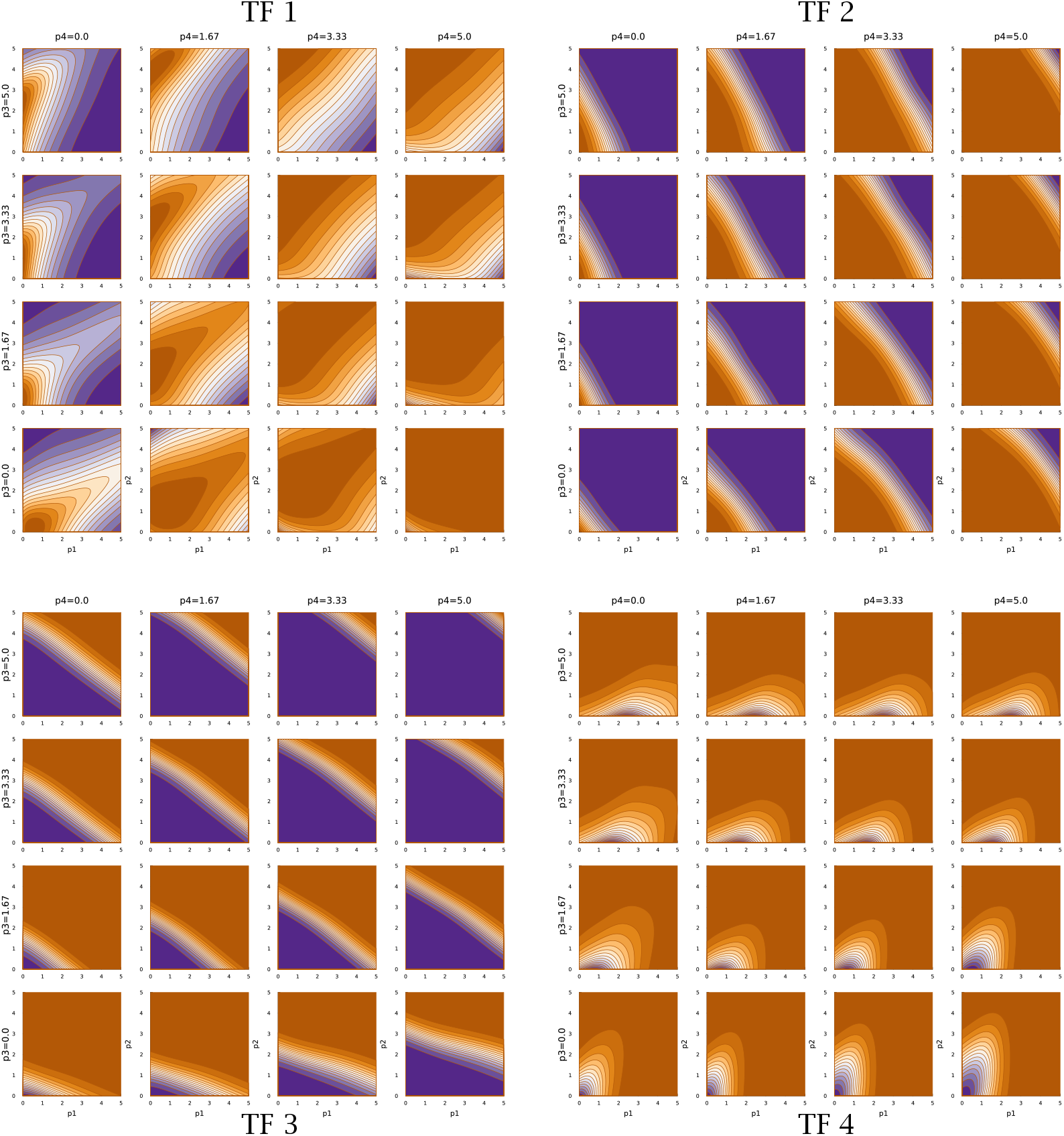
TF network input-output function for run sde–4_2_t4.

**Figure 7:**
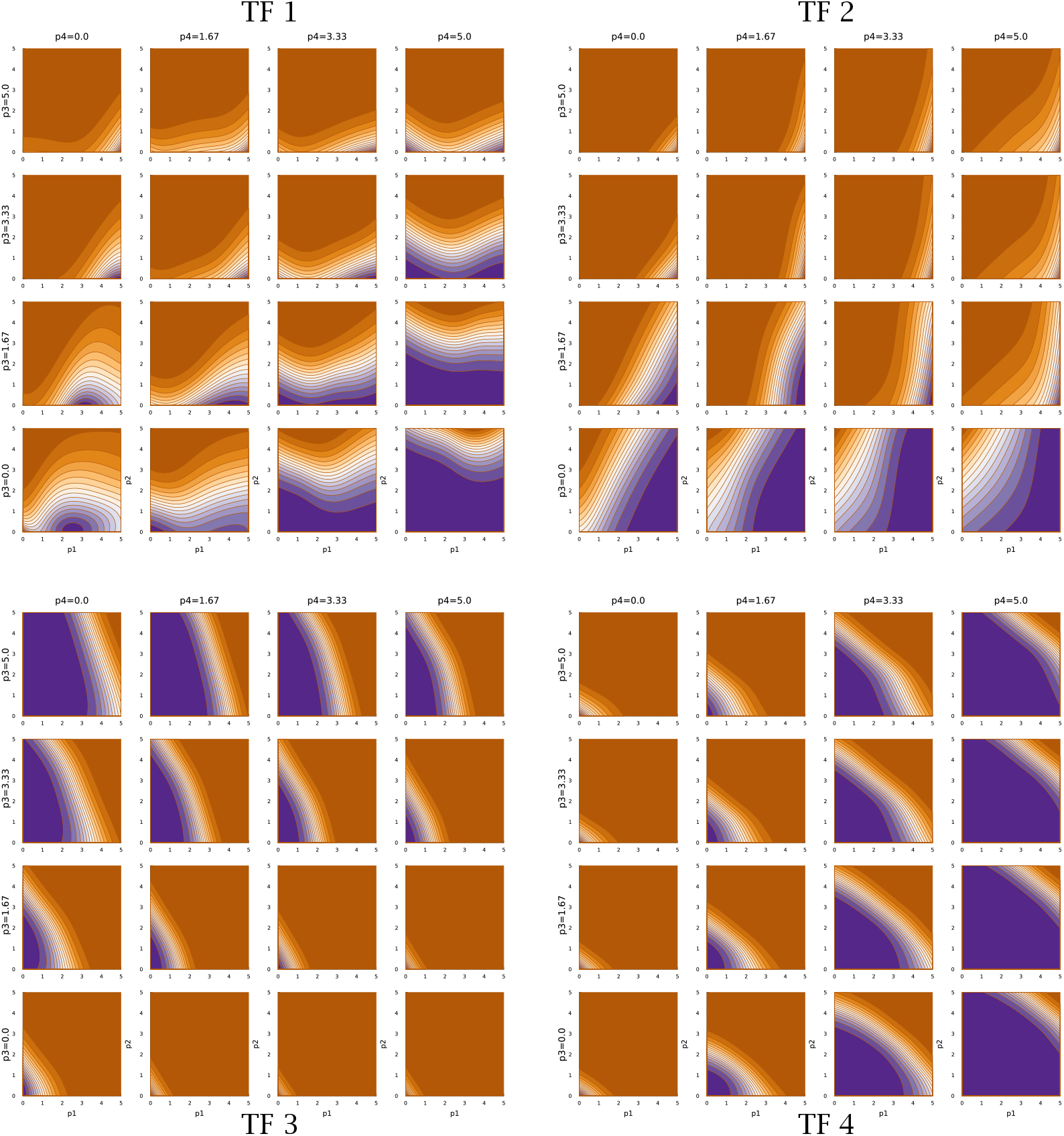
TF network input-output function for run sde–4_8_t4.

Overall, Figs. 5–7 demonstrate that TF networks can produce good performance in different ways. Finding the necessary and sufficient characteristics of the TF networks remains an open problem.

## Discussion

How do cellular regulatory networks solve the challenges of life? Ultimately, that is an empirical question that must be answered by observational and experimental study. However, it may be a difficult question to answer without some conceptual guidance.

Suppose, for example, that the network connections regulating cellular response are dense, deep, and complex. It might be that the network is overparameterized. In other words, the dimensionality of the molecular parameters controlling cellular response may be greater than the dimensionality of the challenges posed by the environment.

Computational deep learning models are typically highly overparameterized artificial neural networks (Radhakrishnan et al., 2020; Poggio et al., 2020; Bartlett et al., 2020). It turns out that overparameterized networks are particularly good at solving complex problems. And, for such overparameterized networks, it is particularly difficult to figure out how the network actually succeeds at solving its challenge (Amey et al., 2021).

One might find some network nodes that work together to achieve a particular component of the broader solution. For example, some nodes might together identify an environmental state or trigger a particular response that feeds into other parts of the network. But, overall, the relations between the individual network components and the response of the network to particular inputs typically remains opaque.

In the same way, it may be difficult to build up an understanding of cellular function by starting with small molecular subnetworks and then expanding analysis to increasingly broader subcomponents. If so, then understanding in theory how such networks might solve problems may provide some guidance.

For example, what are the variety of ways in which cells might buffer against stochastic perturbations to circadian periodicity? How might cells entrain their rhythm to erratic external signals? In overparameterized networks, there is rarely just one solution. Theory can help to identify the range of possible solutions, which could then guide where to look within the complexity of actual cellular regulation.

This article provides an early step on the path to developing theoretical approaches. By using artificial neural networks for the transcription factor network, the computational models can explore the variety of potential solutions to life’s challenges. This article contributes by showing how relatively simple computer code can discover potential solutions.

## Acknowledgments

The Donald Bren Foundation, National Science Foundation grant DEB-1939423, and DoD grant W911NF2010227 support my research.

## Data availability statement

All computer code, parameters and output used to generate the figures are available on Zenodo (Frank, 2022a).

